# In silico labeling enables kinetic myelination assay in brightfield

**DOI:** 10.1101/2022.09.11.507500

**Authors:** Jian Fang, Eun Yeong Bergsdorf, Vincent Unterreiner, Agustina La Greca, Oleksandr Dergai, Isabelle Claerr, Ngoc-Hong Luong-Nguyen, Inga Galuba, Ioannis Moutsatsos, Shinji Hatakeyama, Paul Groot-Kormelink, Fanning Zeng, Xian Zhang

## Abstract

Recent advances with deep neural networks have shown the feasibility of acquiring brightfield images with transmitted light and applying in-silico labeling to predict fluorescent images. We have developed a novel in-silico labeling method based on a generative adversarial network and outperforms the state-of-the-art Unet method in generating realistic fluorescent images and quantitatively recapitulating real staining signals, as demonstrated in a complex co-culture myelination assay. Furthermore, we have performed the assay in live mode with multiple kinetic points, applied in-silico labeling to predict fluorescent images from brightfield and quantified the kinetic phenotypic changes. Thus, the proposed approach provides a potential tool to study the kinetics of cellular phenotypic changes with brightfield imaging.

## Introduction

Myelin sheath is the lipidic structure that concentrically wraps around the axons. In the peripheral nervous system, myelin is formed by the differentiation of the plasma membrane of Schwann cells^1^. It not only allows the rapid saltatory conduction of the action potentials, but also confers protection and nutritional support to axons. Exposure to various factors such as autoimmune insults, trauma or injuries could trigger demyelination, and eventually neurodegeneration. Demyelinating diseases are of clinical importance, including a spectrum of disorders, such as multiple sclerosis and Charcot-Marie-Tooth disease and currently there is no available cure^2^.

*In vivo* models are valuable to assess the molecular mechanisms of demyelination / remyelination, and to validate the effect of pharmacological agents^3^, but have the disadvantages of high complexity and limited throughput. In comparison, an *in vitro* myelination culture assay can be miniaturized and automated and is suited for high throughput profiling and screening of genetic or chemical perturbations.

Generally, primary rodent dorsal root ganglion (DRG) neurons and Schwann cells need to be co-cultured for a few weeks before myelinated axons could be observed, and it is possible to maintain such co-cultures for a much-extended period. The extent of myelination can be assessed by immunostaining with various myelination markers. While *in vitro* myelination takes time to form, immunostaining is an endpoint measurement, so that kinetic information of the myelination process cannot be captured. Also, immunostaining methods are labor and resource intensive as well as error prone. Therefore, there is a need to develop a new method to monitor *in vitro* myelination in live cells.

As the myelinated axons in the co-culture assay are visible with high magnification under transmitted light, we hypothesize that it is possible to establish the assay with brightfield imaging and bypass the challenges of immunostaining. The low signal to noise ratio of brightfield images is generally prohibitive for conventional quantitative image analysis. However, in silico labeling approach has been recently proposed where deep neural network models are trained to map from brightfield images to fluorescent images^4,5^, and thus enables a path to acquire brightfield images, leverage in silico labeling to predict fluorescent signals, and apply conventional quantification on fluorescent images^6–9^.

In this paper, we describe a novel in silico labeling approach, B2FGAN (Brightfield to Fluorescent Generative Adversarial Network), based on the pix2pix generative adversarial network^10^, and demonstrate its application in the myelination assay. The network was trained with 300 images from 60 wells of a 96-well microtiter plate and achieved a high accuracy in terms of predicting the pixel intensity value and the myelination area readout shown by cross validation. Impressively the B2FGAN model successfully predicted the myelin signal based on brightfield imaging even when the experimental fluorescent-labeled staining failed. Furthermore, B2FGAN was applied to an independent data set where brightfield images were acquired weekly for up to 7 weeks and captured kinetic phenotypic changes due to genetic perturbations. Thus, the proposed approach enables the brightfield myelination assay in live mode and provides a powerful tool for investigating demyelinating diseases.

## Results

### Assay setup and B2FGAN workflow

We co-cultured primary rat Schwann cells and DRG neurons for up to seven weeks. The occurrence of in vitro myelination was confirmed by immunostaining with the myelination marker at the end of the experiments (see Methods for details). To develop an in silico labeling method of myelination, we built the B2FGAN workflow (Figure 1a) upon a conditional generative adversarial network (cGAN), named pix2pixHD^10^, for predicting fluorescent labels from brightfield images. The workflow contained a training phase to train the cGAN with paired brightfield and fluorescent-labeled images, and an inference phase to predict corresponding fluorescent signals based on brightfield images.

**Figure 1.**
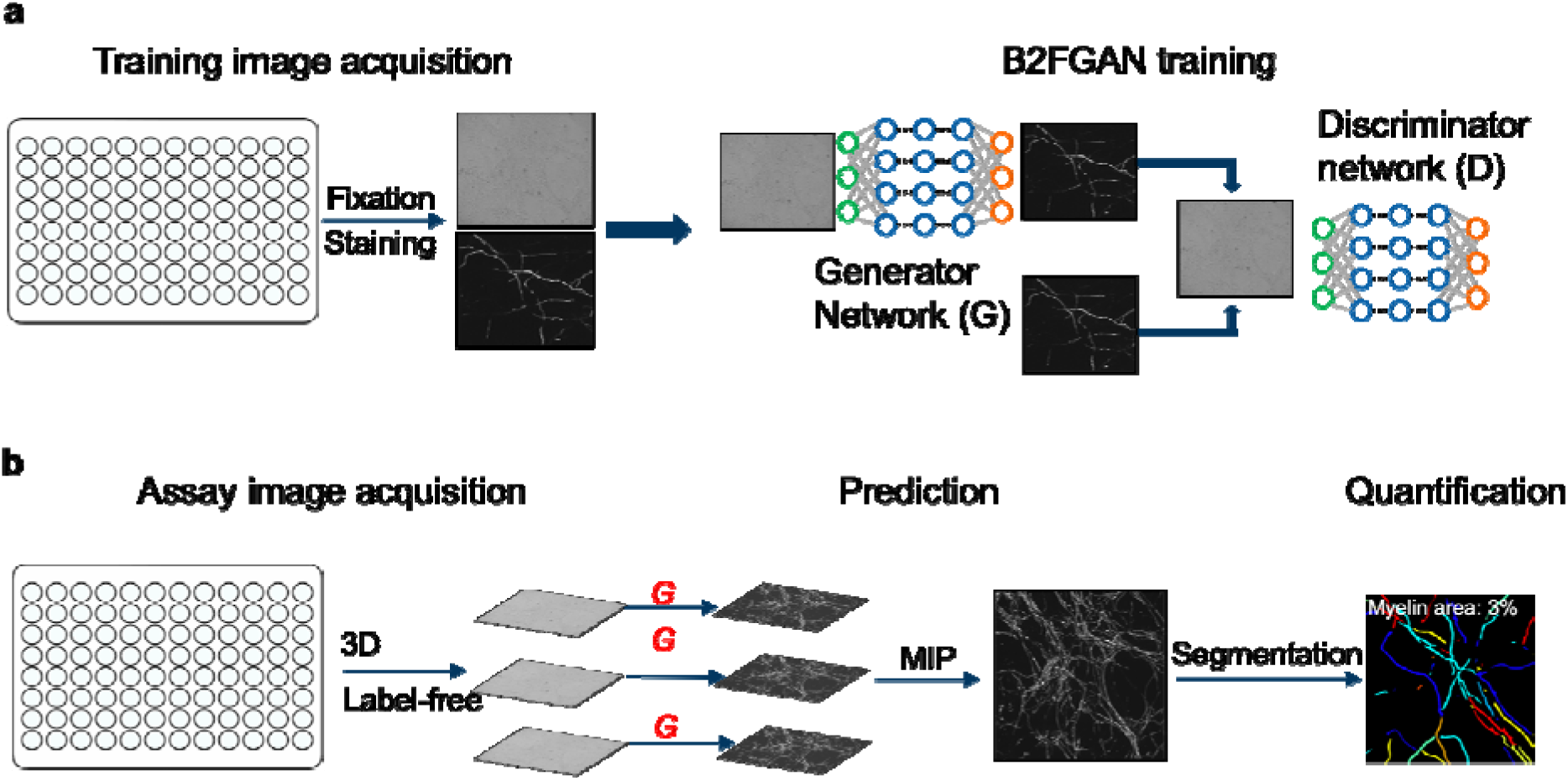
Overall workflow to train the in-silico myelin machine learning model and myelin signal inference. a) Paired brightfield and fluorescent images are collected from 96 well plates with fixed and MBP stained wells. The paired channels are fed into the B2FGAN to learn the generator G that can infer myelin staining from brightfield. b) Given a new brightfield image in 3D, the trained generator predicts each z-stack individually, which are summarized to a single image by maximum intensity projection (MIP), which is subsequently segmented, and the myelin area is measured and reported.

The goal of the training was to learn a mapping from the brightfield channel to the myelin staining channel. To this end, we set up the DRG and Schwann cell co-culture assay, stained with fluorescent-labeled MBP (myelin basic protein), and acquired images from both brightfield and fluorescent channels (details in Methods). Brightfield and myelin-stained images were collected at 20x magnification, with 5 fields of view (FoV) per well. Since the myelin structure is not flat in the plate, 8 z-stacks (with interval of 4 μm) per FoV were captured to account for 3D cellular structure. In addition, the microscope was alternating between the brightfield and myelin channels before moving to the next well, to assure maximum pixel alignment of the two channels (see Methods). The paired brightfield and fluorescent images were then fed into pix2pixHD^10^ for training. The model had two networks, a generator G to generate myelin predictions from brightfield input, and a discriminator D to distinguish real staining and fake generation given the brightfield input. The objective of the GAN training was to learn the data distribution of interest by increasing the error rate of the discriminator, and finally to generate realistic predictions.

During the inference phase (Figure 1b), we took only brightfield images without fixation nor fluorescent-labeled antibody staining. Each z-stack brightfield image was fed into the trained B2FGAN to predict the corresponding myelin signal. The 3D z-stacks of predicted myelin images were subsequently flattened by maximum intensity project (MIP), and the conventional data analysis pipeline segmented the myelin objects and calculated the portion of myelin, i.e., area of myelin divided by the area of whole image, as the readout.

### Method benchmark and validation

With the paired brightfield and myelin images from the 60 wells, we performed 10-fold cross validation with each fold containing 54 wells as training data and 6 wells as test data. We applied the proposed B2FGAN method as well as the previously published Unet^4^ as a baseline method.

Example predictions of the myelin signals are shown in Figure 2a. When compared with the ground truth, both Unet and B2FGAN methods make plausible predictions from brightfield. While the Unet predictions seem blurry, visual inspections suggest that B2FGAN predictions recover more details such as defined boundaries and high variations within the objects, and are more realistic, with sheath-like appearance.

**Figure 2.**
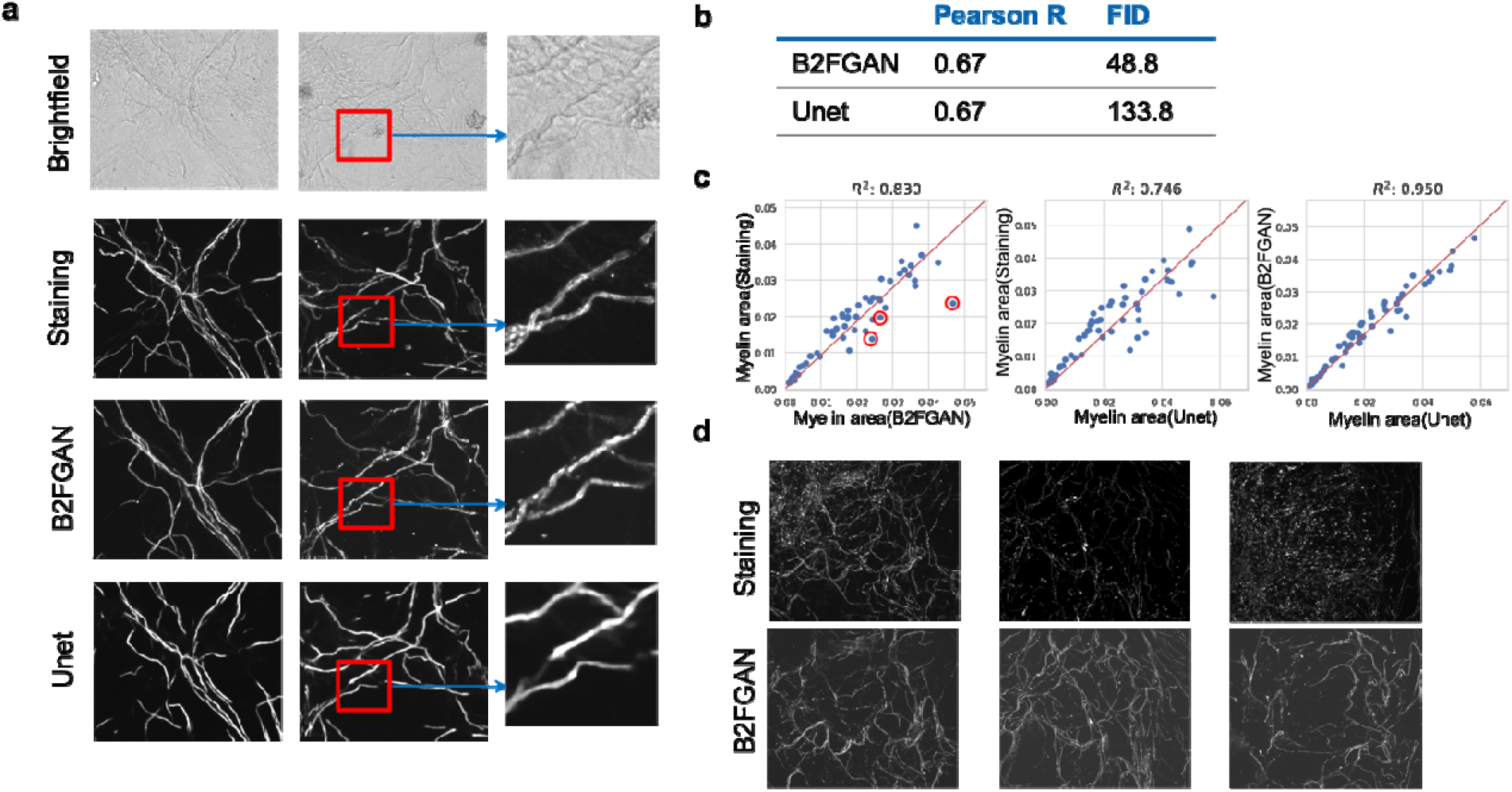
Benchmarking prediction against real staining. a) Example images. From top to bottom are brightfield channel, stained channel, in-silico predictions from B2FGAN and Unet, respectively. The first two columns are two independent examples, and the third column is a zoom-in of the highlighted region in the second column. b) Pearson correlation coefficient of the pixel intensity and Frechet inception distance of image features between in-silico methods (B2FGAN and Unet) and the real staining. c) Correlation of myelination ratio between in silico predictions and stained images. Each dot in the plot corresponds to the averaged myelination area within a well. d) Representative images corresponding to the three points highlighted in c.

To quantitatively evaluate the in-silico labeling methods, we calculated the Pearson’s correlation (*ρ*) between predicted and true pixel intensity, where both B2FGAN and Unet reached a good *ρ* value of 0.67, indicating well recovery of the overall myelin staining at the pixel level (Figure 2b). We also calculated the Frechet inception distance (FID)^11^ of image features between predicted and ground truth images, which evaluates how realistic a prediction is and a lower score is better. B2FGAN achieved a convincingly lower FID than Unet (48.8 vs 133.8, Figure 2b), in agreement with our visual inspections that B2FGAN could generate more realistic myelin signals. Finally, the myelin area values derived from the predictions were compared with those from real staining. As shown in Figure 3c, the *r*^2^ is 0.83 for B2FGAN and 0.75 for Unet. It is interesting to note that *r*^2^ between the two in-silico labeling methods is much higher (0.95), showing that the two methods agree more with each other than with the experimental ground truth. Indeed, close examinations of the outliers (Figure 2d) discovered these wells suffered from inconsistent myelin staining, resulting in artefacts and underestimated myelination. In contrast, the brightfield images were intact and thus the predictions from brightfield showed clearer myelin signal and more accurate quantification.

**Figure 3.**
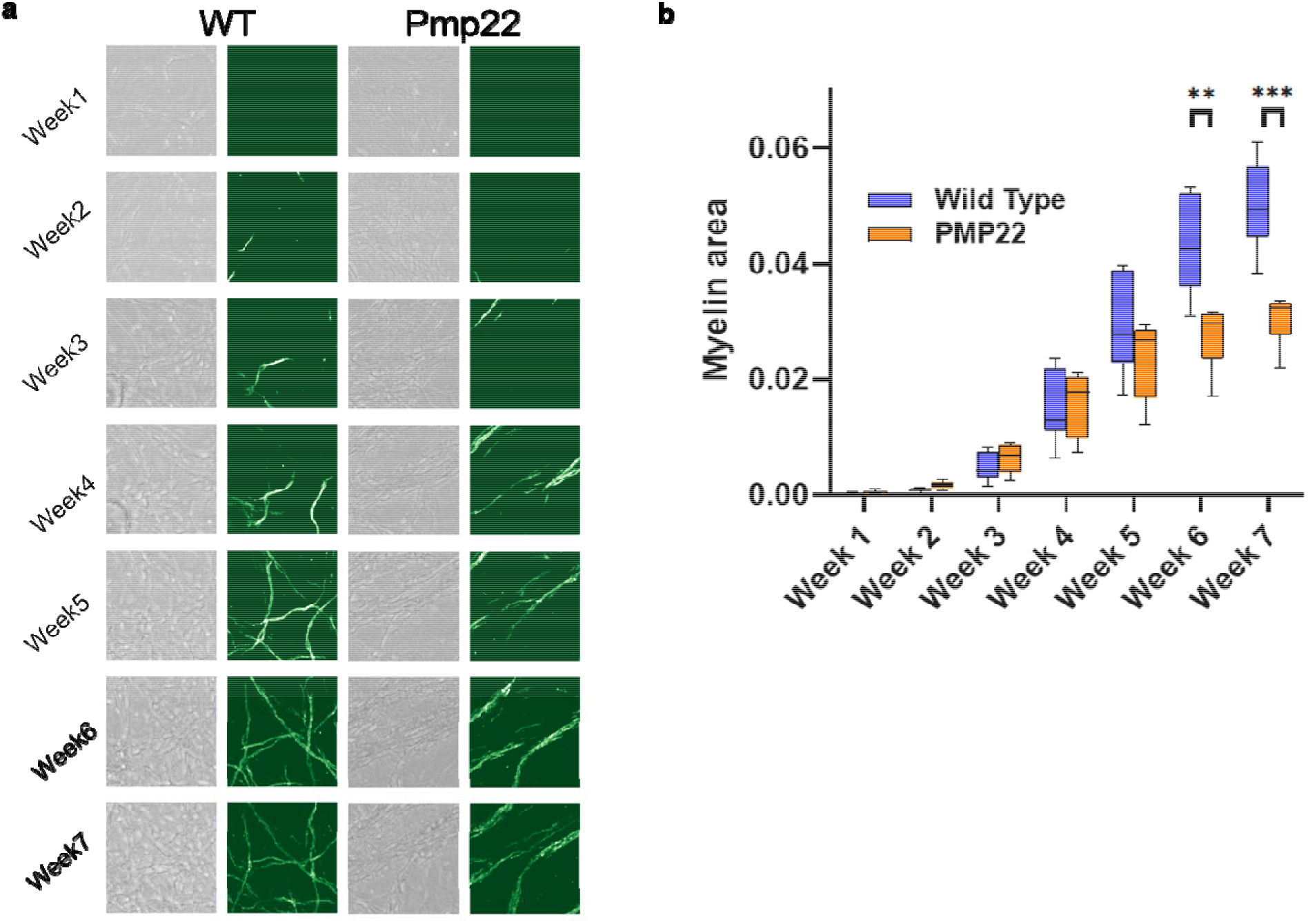
Kinetic assay prediction and analysis results. a) Representative images showing the kinetic of myelin signal during seven weeks. Left, wildtype and right, Pmp22-transgenic cultures. b) Boxplots of myelin area comparing WT and Pmp22-trangenic, from week 1 to week 7.

### Live cell assay and kinetic phenotype

We hypothesize that although the B2FGAN model is trained over images from fixed cells, it can apply to live cell brightfield images and thus enables the assay in live mode and monitoring kinetic myelination processes. To test this, we established the co-culture assay using Schwann cells isolated from wildtype or Pmp22-transgenic rats^12^, collected 3D brightfield images weekly for seven weeks, and applied the previously trained B2FGAN model and quantification as in Figure 1b. Figure 3a shows representative live brightfield images from the same FoV, together with the myelin prediction from B2FGAN. Clear and consistent growth of myelin was observed from the predictions, for both wildtype and Pmp22 cultures.

We further quantified the myelin area readouts across time points between the two genotypes (Figure 3b). Incremental myelin growth was observed for both WT and Pmp22 cultures, but more pronounced in WT. By fitting a four-parameter logistic regression to both genotypes, we had Bayesian information criterion (BIC) (−507.5 for WT and −554.1 for Pmp22) to be higher than linear model (−478.7 for WT and −547.3 for Pmp22) and more than 20% higher than the null model (−341.1 for WT and −410.4 for Pmp22), indicating a logistic behavior of the kinetics. In addition, significantly decreased myelin growth is observed in cultures from Pmp22-transgenic rats (p<0.01, Wilcoxon rank-sum test, Bonferroni correction) at weeks 6 and 7, consistent with the published data that overexpression of Pmp22 is associated with hypomyelination^12,13^.

## Discussion

The field of cellular microscopy analysis has seen substantial progress as diverse neural network based methods are being successfully developed and applied. While most methods aim for the tasks of object segmentation^14,15^ and phenotype classification or clustering^16,17^, two recent studies^4,5^ show the promise of a novel in-silico labeling task, which predicts fluorescent signals from brightfield images. Inspired by the in-silico labeling publications, we have developed the B2FGAN method based on a specific generative adversarial network (GAN) termed pix2pixHD^10^, yielding more realistic myelin images than conventional methods such as Unet. In the realm of computer vision, GAN enables a wide variety of applications, such as image synthesis^18^, image super-resolution^19^, modality translation^20^, with known advantages in producing realistic images than other competitive methods. In contrast, previous work^21^ has found that Unet with *L*_1_ loss may lead to blurred predictions. In the current study, since myelin structures have distinct boundaries and large perimeter to area ratio, the fine details can be readily observed in the B2FGAN predicted images but lost from Unet predictions.

Quantitatively we have evaluated B2FGAN and Unet predictions with three metrics against the ground truth, pixel intensity correlation, myelin area correlation and Frechet inception distance (FID). While the pixel intensity correlation values are the same between the two methods, B2FGAN has achieved better performances in both myelin area correlation and FID. We consider these two metrics are more biologically relevant as the former captures the phenotypic changes and the latter corresponds to biologists′ visual inspection, and thus recommend them for further in-silico labeling method benchmarking studies.

Interestingly, when compared to the ground truth, FID of B2FGAN is much lower than that of Unet, but pixel closeness of these two methods is very similar. Meanwhile, the correlation of the pixel intensity is also much lower than the correlation of the object area, which is indeed more important for capturing the biological changes. Both comparisons inferred that pixel level comparison, which is more often used in previous works^4,5^, would not be enough to evaluate the in-silico labeling approach. This is also supported by another related work^6^, in which morphological feature-level and profiler-level evaluation was added to their in-silico labeling cell painting assay.

After trained with brightfield and fluorescent images from fixed cells, B2FGAN can apply successfully to brightfield images from live cells, which enables the myelination assay to be run in a kinetic mode and be quantified as if it was an end-point assay with fluorescent staining. We show that the live myelination assay with B2FGAN can capture the kinetic phenotypic changes caused by PMP-transgenic cells.

We trained B2FGAN in 2D and applied MIP in the inference stage, instead of training with dense z-stacks^4^, and have achieved comparable or better pixel intensity and assay readout correlation (Figure 2b). Furthermore, we show that the number of z-stacks can be reduced from 32 to 8 without compromising prediction quality (*r*^2^ > 0.98, Supplementary Fig. 1). However, with 1 z-stack, we observe significant quality reduction (*r*^2^ < 0.85), supporting the use of 3D imaging for the myelination assay. Altogether B2FGAN greatly reduces the experimental and computational cost by only using 8 z-stacks. The optimal number of z-stacks in a different assay however will likely vary depending on the biological phenotype.

The B2FGAN training achieved realistic predictions qualitatively and quantitatively with only 300 images from 60 wells. In fact, different from conventional classification tasks, which have a single global label, B2FGAN is based on a local loss function over each pixel of the 2560×2160 images, which brings much more information to guide the training.

The backend algorithm, pix2pixHD^10^, is an efficient choice for high-resolution cellular images. The training takes about 3 days on a single v100 GPU with various ways of augmentations (see Methods for details). The inference is about 3 minutes per well, i.e., 16 FoV and 8 z-stacks in our live assay, for the B2FGAN to generate the predictions. Therefore, we need about 4.8 hours per 96-well plates for weekly kinetic analysis, which could be further accelerated by utilizing multiple GPUs.

The B2FGAN method is also applicable to other cellular imaging assays and brings the same advantages including no need for staining and enabling live assay mode (data not shown). One can hypothesize further that it generalizes to a task where an “easy-to-capture” channel serves as input and a “hard-to-capture” channel is the desired output. Thus, the in-silico labeling approach in general and the B2FGAN method open many opportunities in computationally augmenting cellular microscopy experiments.

## Acknowledgements

We would like to thank Mark-Anthony Bray for his contribution to the work.

## Methods

### Myelin co-culture

#### DRG culture

Embryonic DRG neurons were collected from rat embryos at E14. Briefly, the spinal cord was removed, and the neural tube opened with microdissection tools. The meninges with DRGs attached were peeled off and dissociated by TryPLE (Gibco, cat. no. 12604013) treatment for 1 hour at 37°C. Homogenous cell suspension was obtained by dissociating the tissue with glass Pasteur pipette and filtered through 40 μm cell strainer (Falcon, cat. no. 352340). Dissociated DRG neurons were cultured in complete DRG culture medium containing neuro basal medium (Gibco, cat. no. 12348017) supplemented with 2% B-27 (Gibco, cat. no. 17504044), 2mM L-glutamine (Gibco, cat. no. 25030081) 100 U/mL penicillin-streptomycin (Gibco, cat. no. 15140122) and 50ng/mL nerve growth factor (Alomone, cat. no. N-100) in poly-D-lysine (Sigma, CAS number 27964-99-4) / laminin (Invitrogen, cat. no. 23017015) coated flasks. Proliferating cells were removed by 10uM AraC (Sigma, CAS number 69-74-9) treatment on day 1 and 3. Purified DRG neurons were harvested by TryPLE (Gibco, cat. no. 12604013) treatment and frozen after one week.

#### Schwann cell culture

Primary Schwann cells were isolated as described previously by Kaewkhaw et al. (2012). Briefly, Sciatic nerves were isolated from adult rats (either wildtype or Pmp22-transgenic), and epineurium stripped. The tissues were teased and digested with 0.05% collagenase type IV (Worthington Enzymes, cat. no. LS004188) digestion at 37°C for 2 hours. The cells were maintained in poly-D-lysine coated flasks within Schwann cell growth medium containing DMEM (L-Valine replaced by D-Valine, Bioconcept, cat. no. 1-26S144-I), 2% B-27 (Gibco, cat. no. 17504044), 2 mM L-glutamine (Gibco, cat. no. 25030081), 1 mM sodium pyruvate (Gibco, cat. no. 11360039), 10 mM HEPES (Gibco, cat. no. 15630056), 100 U/mL penicillin-streptomycin (Gibco, cat. no. 15140122) and 10% fetal bovine serum (Gibco, cat. no.16140071) supplemented with 5 μM forskolin (Sigma, CAS no. 66575-29-9). Once the culture reached to above 80% confluency, Schwann cells were dispatched by DispaseII (Roche, cat. no. 4942078001) digestion and can be sub-cultured for up to 10 passages. The purities of Schwann cell were at least 95%, judged by immunostaining with Schwann cell markers.

### DRG Schwann cell co-culture

Cryopreserved DRG neurons were thawed quickly by placing them in into a 37°C water bath. The DRG neurons were resuspended with complete DRG culture medium. A 96-well plate (PerkinElmer, cat. no. 6005182) was coated with 4% matrigel (Corning, cat. no. 354277) diluted with X-vivo 10 medium (Lonza, cat. no. BE04-3800) for minimum 12 hours at 4°C before DRG neuron seeding. The DRG neurons were seeded at a density of 2,900 cells/well. One hour after DRG neuron seeding, cryopreserved Schwann cells were defrosted in a similar way to the DRG neurons. The Schwann cells were resuspended with Schwann cell growth medium and seeded at density of 32,000 cells/well. The medium was changed to 100% complete DRG culture medium supplemented with 5 μg/mL ascorbic acid (Waco, CAS no. 1713265-25-8) at the 4th day after cell seeding. Half of culture medium was exchanged to fresh DRG culture medium throughout the culture period.

### Immunostaining

Immunostaining was performed according to standard protocols. Cells grown on 96 well plates were fixed and permeabilized for 20 mins at 4°C with Fixation/Permeabilization solution (BD Biosciences, cat. no. 554714). Cells were washed 3 times in PBS, then blocked in PBS containing 5% donkey serum (Sigma, cat. no. D9663), 1 % bovine serum albumin (BSA, Jackson, cat. no. 001-000-162) and 10% Perm/Wash buffer (BD Biosciences, cat. no. 554714) at RT for 1 hour. For antibody staining, cells were incubated overnight at 4°C with 1:100 Myelin Basic Protein antibody (Bio Rad, cat. no. MCA409S), washed three times with PBS, then incubated with 1:1000 Alexa Fluor-conjugated secondary antibodies (ThermoFisher) for 1 hour at RT. The Nuclei were visualized by counterstaining with NucBlue (Invitrogen, cat. no. R37606). Finally, the cells were rinsed a further 3 times in PBS (Gibco. cat. no. 14190-094) before reading the plate on Yokogawa CV7000.

### Generation of the training set images

After fixation and staining, the plates containing the cells (PerkinElmer, ViewPlate-96, cat# 6005182) were imaged on a CV7000 automated microscope (Yokogawa). In each well, 16 fields of view were pictured. For each field of view, the brightfield and fluorescent images were acquired sequentially using the same confocal light path to ensure the best pixel alignment between the two acquisition channels. Images were taken using a 20X magnification objective with a 0.75-NA (Olympus; uplsapo20x) on 8 different z-slices with an inter-z-slice interval of 4 μm. The binning factor of the camera was set to one, the image X-Y resolution was 2560×2160 pixels and the image pixel size was 0.325 μm/pixel. The acquisition settings for each channel were the following: Channel 1 (brightfield): Lamp, 100 ms exposure, emission filter 525/50 nm; Channel 2 (MBP staining): 488nm laser, 250 ms exposure, emission filter 525/50 nm; Channel 3 (Nuclei staining): 365nm laser, 750 ms exposure, emission filter 445/45 nm.

### Live cell imaging

For live cell measurement, cells were imaged using the brightfield confocal light path of the CV7000. Cells were kept at 37°C and 5%CO2 in a humidified environment inside the device. In each well, 16 fields of view were imaged using a 20X magnification objective with a 0.75-NA (Olympus; uplsapo20x) on 8 different z-slices with an inter-z-slice interval of 4 μm. The binning factor of the camera was set to one, the image X-Y resolution was 2560×2160 pixels and the image pixel size was 0.325 μm/pixel. The acquisition settings were the following: Channel 1 (brightfield): Lamp, 100 ms exposure, emission filter 525/50 nm.

### Model architecture

We employed the original pix2pixHD^10^ architecture to train the mapping from the brightfield channel to the corresponding myelin channel. The pix2pixHD is a conditional GAN framework, optimized for high-resolution images and consists of a coarse-to-fine generator and a multi-scale discriminator. The generator takes the brightfield image as input, and outputs a fake staining image with the same size. The discriminator takes the pair of brightfield and stained images as input and classifies whether it is real or fake. The two networks are optimized via a minimax game to model the conditional distribution of real myelin staining over the brightfield image. Conditional GAN was well known to generate more realistic images than conventional methods, e.g., Unet. With improved networks and adversarial loss, pix2pixHD can deal with very high-resolution data, e.g., 2560×2160 in this work.

Unet^4^ is also included for the comparison. Unet is a convolutional neural network which has proven very effective in many imaging tasks^22^. U-Nets involve several convolutions in a contracting path for encoding, and an expansive path for decoding to learn the feature relationships. The network architecture used in this paper is given in supplementary Fig 2.

### Model training

Both models were trained over 200 epochs on a single Nvidia v100 GPU using Pytorch. The input batch size is 1 pair of 2D BF and myelin slice for pix2pixHD and 4 pairs of 3D BF and myelin images for Unet. Instead of preprocessing, within each batch, on-the-fly augmentation was applied, including random cropping, resizing and perturbations over normalization. More specifically, denote *Q(p)* be the th quantile of the pixel intensity of an image, we estimate

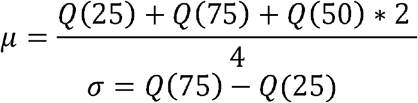

Then the image is centered by removing *r*_1_*μ* and normalized by dividing *r*_1_*σ*, where *r*_1_ and *r*_2_ are sampled from *U*[0.9,1.1]. After normalization, the image was resized by a factor randomly drawn from [0.9,1.1]. Finally, a random patch with size 1024×1024 is cropped from the resized image.

For pix2pixHD, we adopt the default parameters for training. And to train Unet, we minimized over huber loss, and used ADAM with learning rate of 1e-3 as the optimizer.

### Model inference

For both B2FGAN and Unet, the full size brightfield image was normalized to be the input. The output of the generator is mapping back to 16-bit images by inverting the scaling and shifting using the mean *μ* and *σ* from the training data.

### Cellprofiler pipeline for myelin quantification

A CellProfiler^23^ pipeline was utilized to segment myelin and estimate the myelin area. The pipeline mainly consists of three modules, subsequently to enhance neurites features, to identify primary myelin objects, and to estimate the area occupied by the myelin. The pipeline with detailed parameters can be found in the supplementary data.

### Evaluation metrics

To evaluate the performance of myelin prediction, three different metrics were used, the Pearson’s R over the pixel intensity, the Frechet inception distance (FID) over image features, and *r*^2^ over the myelin area derived from the cellprofiler pipeline.

More specifically, the Pearson’s R measures the linear correlation of pixel intensity between the predicted image and the real staining, defining as

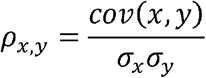

Where *cov(x, y)* is the covariance between *X* and *Y*, *σ*_*x*_ and *σ*_*y*_ are standard deviations of *x* and *y*.

The FID measures the distance between feature vectors of predicted and real staining. The score compared the statistics of the image patches between two groups, with respect to image features derived by pretrained Inception v3 model. The real stained images were first divided into 299×299 patches, which is the input size of the Inception v3 model. Then, a subset of 1709 patches with myelin area (calculated from the CellProfiler pipeline) greater than 5% was picked, to reduce the noisy contribution from too sparse images. The real staining patches, and the corresponding predicted patched were fed into the pre-trained Inception v3 model, and the output of feature layers were kept as *X* and *Y*, the FID then is calculated as

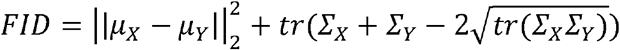

Where *μ* is the mean vector and *Σ* is the covariance matrix.

*r*^2^ was used to measure the linear correlation of assay readout, i.e., myelin area from the CellProfiler pipeline, between the real staining and predictions.

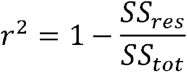

Where *SS*_*tot*_ = *Σ*_*i*_(*y*_*i*_ – *μ*_*y*_)^2^, and *SS*_*res*_ = *Σ*_*i*_(*y*_*i*_ – *x*_*i*_)^2^.

### Kinetic data analysis

To access the behavior of myelin growth, Bayesian information criteria (BIC) was used to determine if the myelin area as a function of time was best fit by a linear model, a four-parameter logistic model, or a “null” model (no change over time).

