## Supplementary figures for "In silico labeling enables kinetic myelination assay in brightfield"

**Figure 1**

Effects of z-stack interval to the myelin quantification, evaluated over the real staining. a) Scatter plot comparing the myelin area from a single z-stack and the maximum intensity projection (MIP) from 32 z-stacks. b) Scatter plot comparing MIP from 4x down-sampling (8 z-stacks) and 32 z-stacks. c) Correlation of myelin area as a function of down-sampling factors.

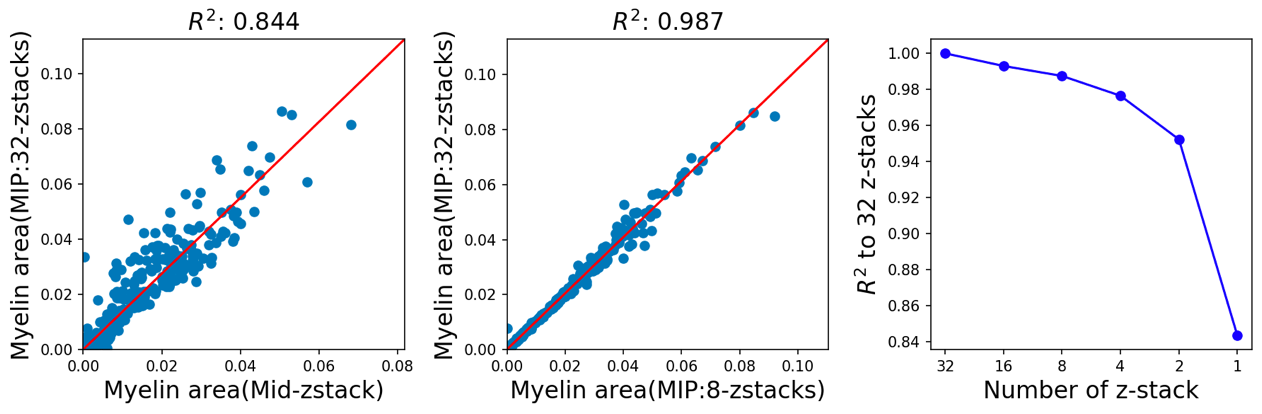

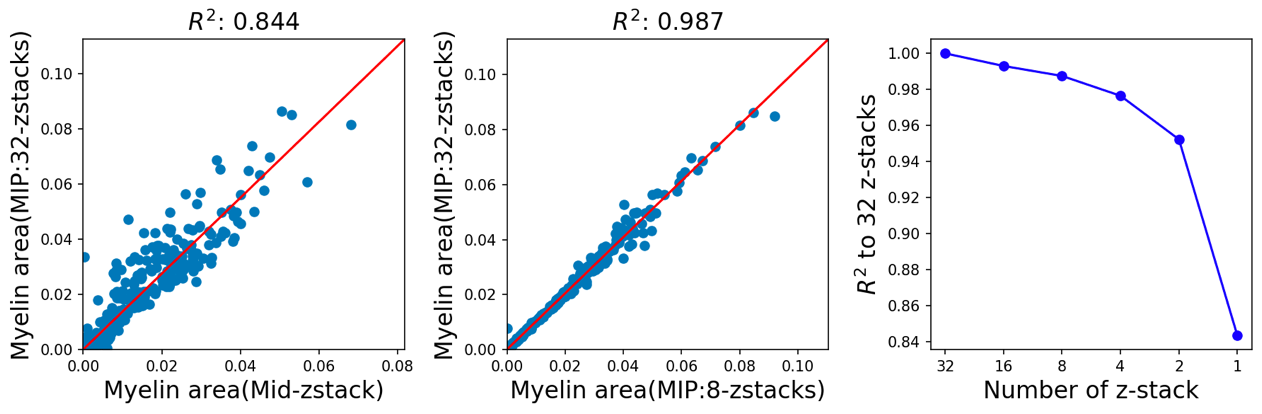

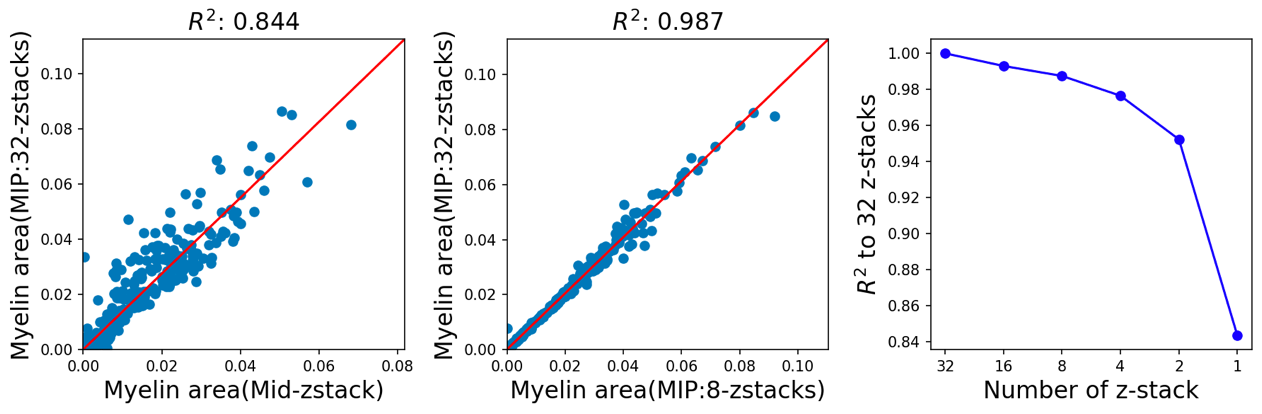

**Figure 2**

Network architecture for 3D Unet we used as benchmark method in the paper. In conv boxes, the number refers to kernel size (width, height, z), output channel size, padding size. In pooling and up-sampling boxes, the number refers to kernel size.

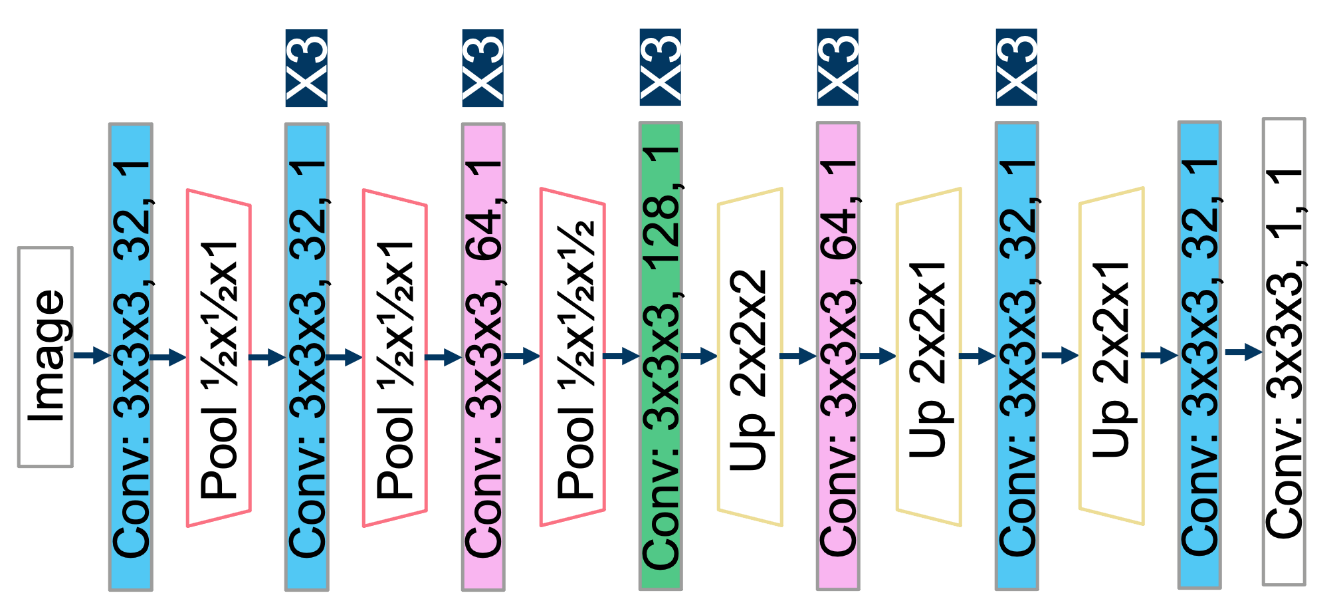
